# FastGT: an alignment-free method for calling common SNVs directly from raw sequencing reads

**DOI:** 10.1101/060822

**Authors:** Fanny-Dhelia Pajuste, Lauris Kaplinski, Märt Möls, Tarmo Puurand, Maarja Lepamets, Maido Remm

## Abstract

We have developed a computational method that counts the frequencies of unique *k*-mers in FASTQ-formatted genome data and uses this information to infer the genotypes of known variants. FastGT can detect the variants in a 30x genome in less than 1 hour using ordinary low-cost server hardware. The overall concordance with the genotypes of two Illumina “Platinum” genomes^1^ is 99.96%, and the concordance with the genotypes of the Illumina HumanOmniExpress is 99.82%. Our method provides *k*-mer database that can be used for the simultaneous genotyping of approximately 30 million single nucleotide variants (SNVs), including >23,000 SNVs from Y chromosome. The source code of FastGT software is available at GitHub (https://github.com/bioinfo-ut/GenomeTester4/).

---

Next-generation sequencing (NGS) technologies are widely used for studying genome variation. Variants in the human genome are typically detected by mapping sequenced reads and then performing genotype calling^2–5^. A standard pipeline requires 40-50 h to process a human genome with 30x coverage from raw sequence data to variant calls on a multi-thread server. Mapping and calling are state-of-the-art processes that require expert users familiar with numerous available software options. It is not surprising that different pipelines generate slightly different genotype calls^6–10^. Fortunately, inconsistent genotype calls are associated with certain genomic regions only^11–13^, whereas genotyping in the remaining 80-90% of the genome is robust and reliable.

The use of *k*-mers (substrings of length *k*) in genome analyses has increased because computers can handle large volumes of sequencing data more efficiently. For example, phylogenetic trees of all known bacteria can be easily built using *k*-mers from their genomic DNA^14–16^. Bacterial strains can be quickly identified from metagenomic data by searching for strain-specific *k*-mers^17–19^. *K*-mers have also been used to correct sequencing errors in raw reads^20–23^. One recent publication has described an alignment-free SNV calling method that is based on counting the frequency of k-mers^24^. This method converts sequences from raw reads into Burrows-Wheeler transform and then calls genotypes by counting using a variable-length unique substring surrounding the variant.

We developed a new method that offers the possibility of directly genotyping known variants from NGS data by counting unique *k*-mers. The method only uses reliable regions of the genome and is approximately 1-2 orders of magnitude faster than traditional mapping-based genotype detection. Thus, it is ideally suited for a fast, preliminary analysis of a subset of markers before the full-scale analysis is finished.

The method is implemented in the C programming language and is available as the FastGT software package. FastGT is currently limited to the calling of previously known genomic variants because specific *k*-mers must be pre-selected for all known alleles. Therefore, it is not a substitute for traditional mapping and variant calling but a complementary method that facilitates certain aspects of NGS-based genome analyses. In fact, FastGT is comparable to a large digital microarray that uses NGS data as an input. Our method is based on three original components: 1) the procedure for the selection of unique *k*-mers, 2) the customized data structure for storing and counting *k*-mers directly from a FASTQ file, and 3) a maximum likelihood method designed specifically for estimating genotypes from *k*-mer counts.

## RESULTS

### Compilation of the database of unique *k*-mer pairs

The crucial component of FastGT is a pre-compiled flat-file database of genomic variants and corresponding *k*-mer pairs that overlap with each variant. Every bi-allelic single nucleotide variant (SNV) position in the genome is covered by *k k*-mer pairs, where pair is formed by *k*-mers corresponding to two alternative alleles (Figure S4). FastGT relies on the assumption that at least a number of these *k*-mer pairs are unique and appear exclusively in this location of the genome; therefore, the occurrence counts of these unique *k*-mer pairs in sequencing data can be used to identify the genotype of this variant in a specific individual.

The database of variants and unique *k*-mers is assembled by identifying all possible *k*-mer pairs for each genomic variant and subjecting them to several steps of filtering. The filtering steps remove variants for which unique *k*-mers are not observed and variants that produce non-canonical genotypes (non-diploid in autosomes and non-haploid in male X and Y chromosomes) in a sequenced test-set of individuals. Filtering of *k*-mers was performed using high coverage NGS data of 50 individuals from Estonian Genome Project (published elsewhere). The filtering steps are described in Methods section and in Supplementary File (Figure S5).

Although one *k*-mer pair is theoretically sufficient for genotyping, mutations occasionally change the genome sequence in the neighborhood of an SNV, effectively preventing the detection of the SNV by a chosen *k*-mer. If the mutation is allele-specific, then the wrong genotype could be easily inferred. Therefore, we use up to three *k*-mer pairs per variant to prevent erroneous calls caused by the occasional loss of *k*-mers because of rare mutations.

In the current study, we compiled a database of all bi-allelic SNVs from dbSNP and tested the ability of FastGT to detect these SNVs with 25-mers. After the filtering steps, 30,238,283 (64%) validated and bi-allelic SNVs remained usable by FastGT. We also used a subset of autosomal SNV markers present on the Illumina HumanOmniExpress microarray for a concordance analysis. In this set, 78% of the autosomal markers from this microarray were usable by FastGT. The number of SNV markers that passed each filtering step is shown in Table S1.

### Algorithm and software for *k*-mer-based genotyping

The genotyping of individuals is executed by reading the raw sequencing reads and counting the frequencies of *k*-mer pairs described in the pre-compiled database of variants using the custom-made software 

~~~
gmer_counter
~~~

 and 

~~~
gmer_caller
~~~

 (Figure 1).

**Figure 1.**
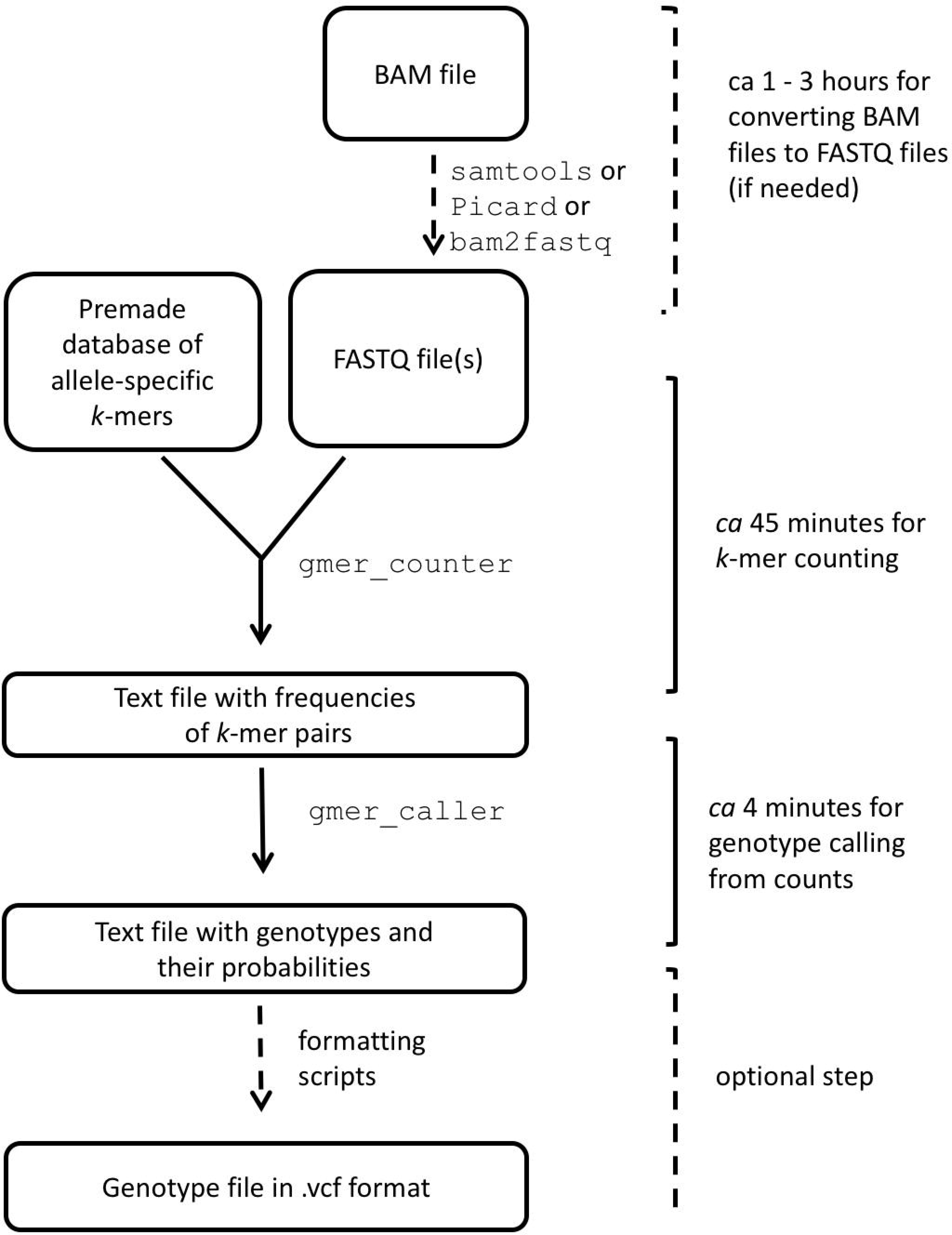
Overall principle of *k*-mer-based genotyping.

The database of genomic variants and corresponding *k*-mers is stored as a text file. The frequencies of *k*-mers listed in the database are counted by 

~~~
gmer_counter
~~~

. It uses a binary data structure, which stores both *k*-mer sequences and their frequencies in computer memory during the counting process. A good compromise between memory consumption and lookup speed is achieved by using adaptive radix tree (see Supplementary File for detailed description of the data structure). The first 14 nucleotides of a *k*-mer form an index into a table of sparse bitwise radix trees that are used for storing the remaining sequence of the *k*-mers. Two bytes per *k*-mer are allocated for storing frequencies. The current implementation of 

~~~
gmer_counter
~~~

 accepts *k*-mers with lengths between 14 and 32 letters. The frequencies of up to three *k*-mer pairs from 

~~~
gmer_counter
~~~

 are saved in a text file that is passed to 

~~~
gmer_caller
~~~

, which infers the genotypes based on *k*-mer frequencies and prints the results to a text file.

### Empirical Bayes’ method for inferring genotypes from *k*-mer counts

~~~
Gmer_caller
~~~

 uses the Empirical Bayes classifier for calling genotypes from *k*-mer frequency data, which assigns the most likely genotype to each variant. Allele frequency distributions are modeled by negative binomial distribution, described by eight parameters (see description in the Supplementary File). The model parameters are estimated separately for each analyzed individual using *k*-mer counts of 100,000 autosomal markers. The model allows us to estimate the most likely copy number for both alleles. Given the observed allele counts, 

~~~
gmer_caller
~~~

 calculates the probability of genotypes by applying the Bayes rule. As we do not require allele copy numbers to sum to 2 we can also call mono-, tri-, or tetra-allelic genotypes (which might correspond to deletions and duplications) in addition to traditional bi-allelic (diploid) genotypes (Figure 2). The model parameters can be saved and re-used in subsequent analyses of the same dataset.

**Figure 2.**
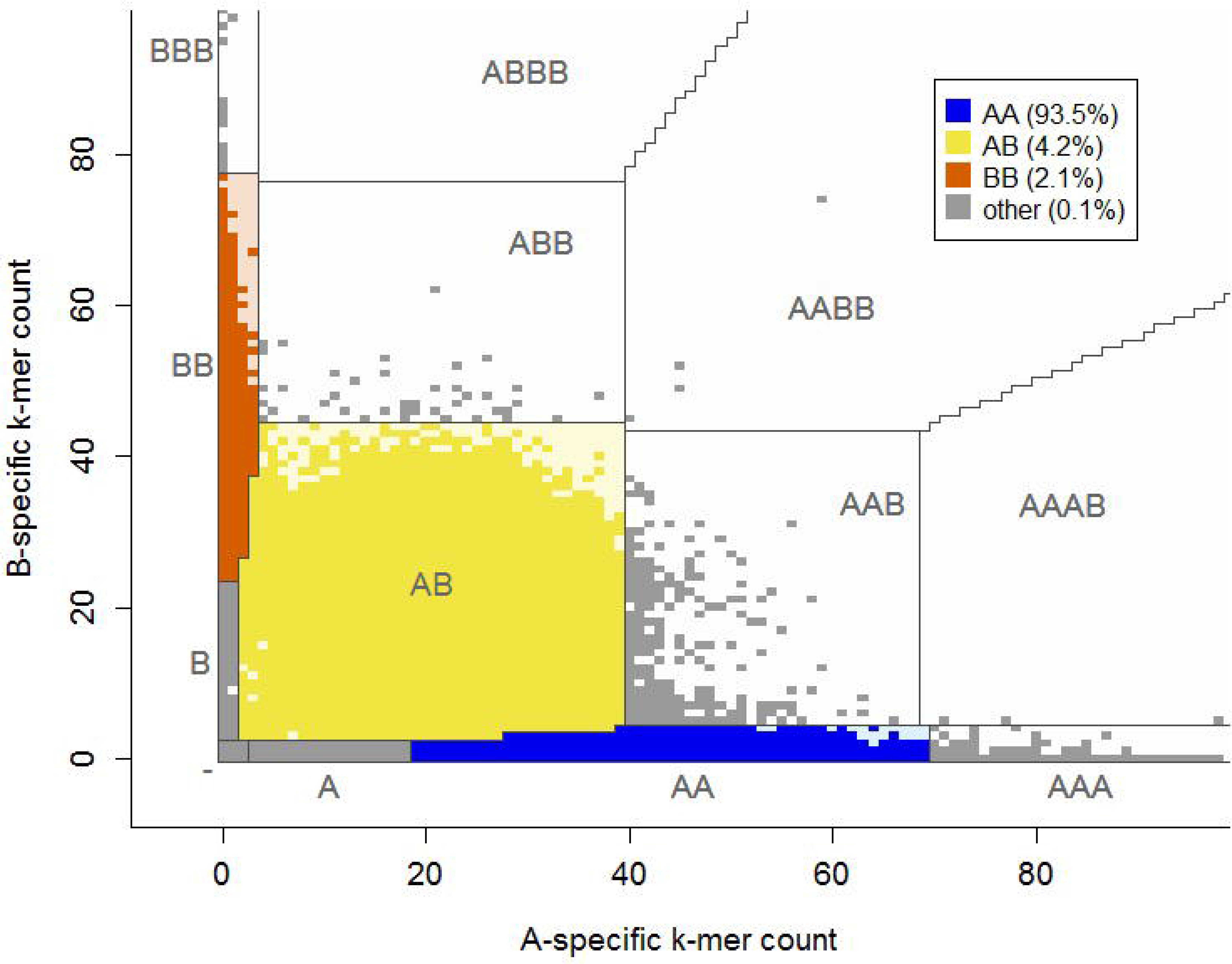
Illustration of genotype calling based on the frequencies of two *k*-mers. The parameters that define boundaries between genotypes are estimated from the *k*-mer frequency data of each individual. By default, only conventional genotypes are reported in the output. “A” denotes the reference allele, and “B” denotes an alternative allele. The estimated depth of coverage (λ) of the individual used in this example was 38.6.

The gender of the individual is determined automatically from the sequencing data using the median frequency of markers from the X chromosome (chrX). If the individual is female, only the autosomal model is used in the calling process and Y chromosome (chrY) markers are not called. For men, an additional haploid model of Bayes’ classifier is trained for calling genotypes from sex chromosomes. Parameters for the haploid model are estimated using 100,000 markers from chrX.

### Assessment of genotype calling accuracy through simulations

In order to test the performance of FastGT, we generated simulated raw sequencing reads from the reference genome and analyzed the ability of the Bayesian classifier of FastGT to reproduce genotypes of the reference genome (see Methods for detailed description of data simulation methods). Throughout this paper we denote A as reference allele and B as alternative allele. In this simulation, the reference genome was assumed to be homozygous in all positions. Thus, the correct genotype for all 30,238,283 tested markers would be AA genotype. The fraction of AA genotypes recovered from simulated reads varied between 98.94% (at 5x coverage) and 99.95% (at 20x coverage). The fraction of uncalled markers was between 0.001% (at 20x coverage) and 1.036% (at 5x coverage). The fraction of AB calls was in range of 0.02% to 0.05% at all coverages. The results are shown in Table 1.

**Table 1.** Genotypes retrieved from the simulated reads generated from the reference genome. “A” denotes the allele from the reference genome, and “B” denotes the alternative allele. “NC” is no-call.

|  |  | Coverage |  |  |  |  |
| --- | --- | --- | --- | --- | --- | --- |
|  |  | 5x | 10x | 20x | 30x | 40x |
| FastGT genotype calls | AA | 28,734,597<br>(98.943%) | 29,025,104<br>(99.943%) | 29,027,872<br>(99.952%) | 29,025,157<br>(99.943%) | 29,011,146<br>(99.895%) |
|  | AB | 6,140<br>(0.021%) | 12,107<br>(0.042%) | 13,567<br>(0.047%) | 11,007<br>(0.038%) | 10,898<br>(0.038%) |
|  | BB | 0<br>(0%) | 0<br>(0%) | 0<br>(0%) | 0<br>(0%) | 0<br>(0%) |
|  | NC | 300,941<br>(1.036%) | 4,467<br>(0.015%) | 239<br>(0.001%) | 5,514<br>(0.019%) | 19,634<br>(0.068%) |

The performance of calling AB and BB genotypes cannot be estimated from the reference genome. We created simulated genomes using genotypes from 5 sequenced individuals, each from different population (Yoruban, Chinese Han, CEPH, Puerto Rican and Estonian). This analysis helped to test the performance of Bayesian classifier (

~~~
gmer_caller
~~~

) on calling the AB and BB variants from real-life data. Secondly, this analysis indicates whether the selection of markers that was done using Estonian individuals introduces any population-specific bias in genotype calling. The sensitivity (fraction of correctly called AB and BB variants) was strongly affected by coverage (61% at 5x coverage, 99.8% at 20x coverage), but remained almost identical for individuals from different populations: 99.7 − 99.8% at 30x coverage (Figure 3). The specificity (fraction of correctly called AA calls) was more uniform over different coverage levels and remained between 99.60% to 99.95%. These results show that our set of 30 million markers is usable for studying different populations without strong bias in sensitivity or specificity.

**Figure 3.**
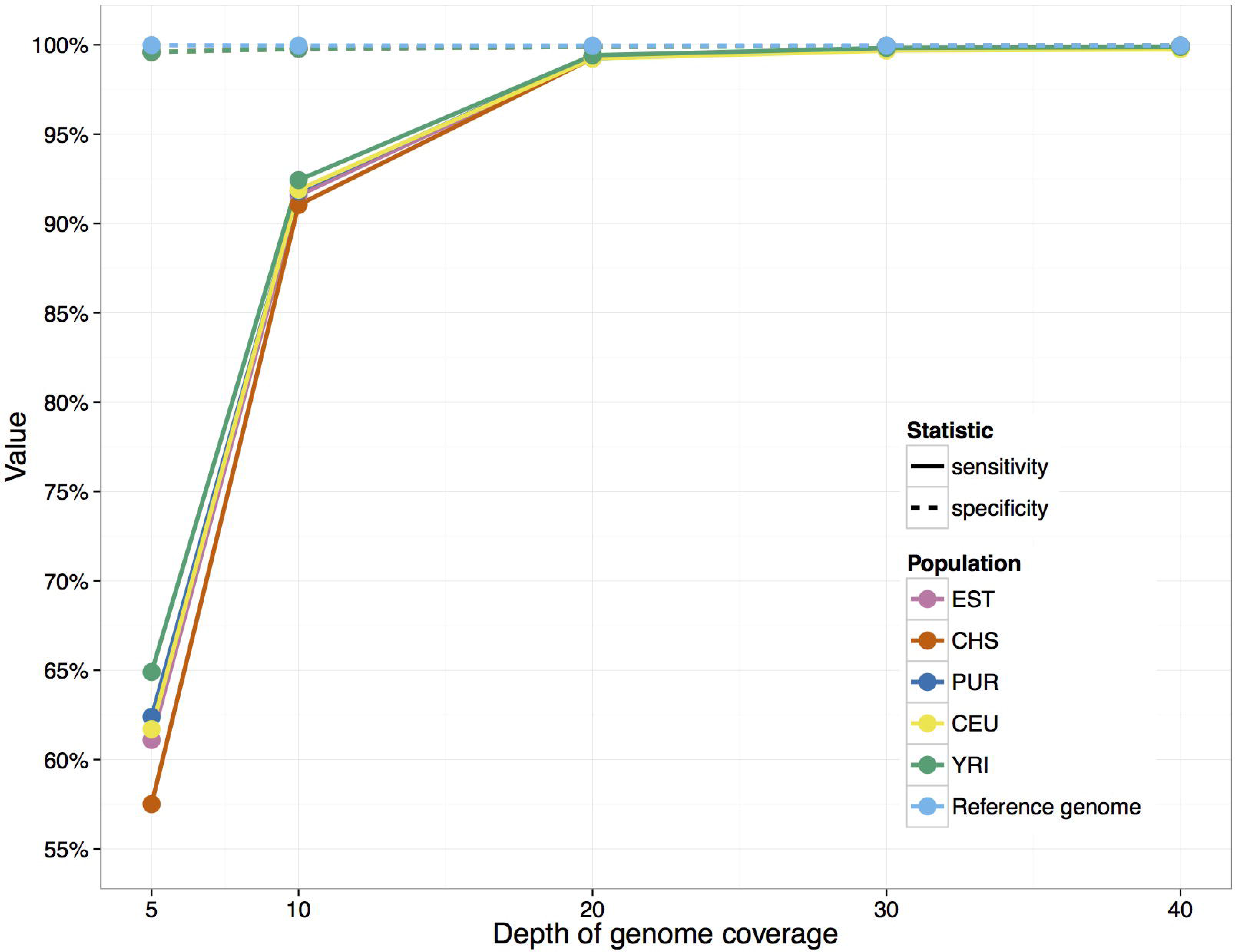
Performance of Bayesian classifier with simulated data. Sensitivity and specificity of calling alternative alleles is shown for reference genome and simulated genome of individuals from 5 different populations. Populations are abbreviated as follows: EST – Estonian; CHS – Southern Han Chinese; PUR – Puerto Rican; CEU – Utah residents with Northern and Western European Ancestry; YRI – Yoruban from Ibadan, Nigeria. Specificities are nearly identical for all individuals and thus their lines are overlapping with the green dotted line.

### Assessment of genotype calling accuracy through concordance analysis

The accuracy of FastGT genotype calls was analyzed by comparing the results to genotypes reported in two Illumina Platinum individuals, NA12877 and NA12878, which were sequenced to 50x coverage. These are high-confidence variant calls derived by considering the inheritance constraints in the pedigree and the concordance of variant calls across different methods^1^. We determined genotypes for 30,238,283 millions of markers from the FastGT database using raw sequencing data from the same individuals and compared them to genotypes shown in the Platinum dataset.

The overall concordance of bi-allelic FastGT genotypes with genotypes from two Platinum genomes is 99.96%. The concordance of the non-reference (AB or BB) calls was 99.93%. The distribution of differences between the two sets for different genotypes is shown in Table 2. All of the genotypes reported in the Platinum datasets were bi-allelic; thus, we included only bi-allelic FastGT genotypes in this comparison. The fraction of uncertain (no-call) genotypes in the FastGT output was 0.24%. The uncertain genotypes are primarily mono-allelic (A) and tri-allelic (AAA) genotypes that might correspond to deletions or insertions in a given region. However, non-canonical genotypes in the default output are not reported, and they are replaced by NC (“no call”). All of the genotypes and/or their likelihoods can be shown in 

~~~
gmer_caller
~~~

 optional output.

**Table 2.** Concordance between the autosomal genotypes of two individuals from the Platinum dataset and bi-allelic FastGT genotypes called from the same individuals. The reference allele is denoted by “A” and the alternative allele is denoted by “B” denotes the alternative allele.

|  |  | Platinum genotype calls |  |  |
| --- | --- | --- | --- | --- |
|  |  | AA | AB | BB |
| FastGT genotype calls | AA | 54,246,425 (93.39%) | 987 (0.00%) | 68 (0.00%) |
|  | AB | 20,041 (0.03%) | 2,427,315 (4.18%) | 1,516 (0.00%) |
|  | BB | 2,261 (0.00%) | 156 (0.00%) | 1,245,902 (2.14%) |
|  | NC | 126,513 (0.22%) | 1,787 (0.00%) | 10,376 (0.02%) |
| concordant (%) |  | 99.96% | 99.95% | 99.87% |

We also compared the genotypes obtained by the FastGT method with the data from the Illumina HumanOmniExpress microarray. We used 504,173 autosomal markers that overlap our whole-genome dataset (Table S2), and the comparison included ten individuals from the Estonian Genome Center for whom both microarray data and Illumina NGS data were available. In these 10 individuals, the concordance between the genotypes from the FastGT method and microarray genotypes was 99.82% (Table 2), and the concordance of non-reference alleles was 99.69%. The fraction of mono-allelic and tri-allelic genotypes (no-call genotypes) in 10 test individuals is rather low (<0.01% of all markers), indicating that our conservative filtering procedure is able to remove most of the error-prone SNVs.

### Markers from Y chromosome

FastGT is able to call genotypes from the Y chromosome (chrY) for 23,832 markers that remain in the whole-genome dataset after all filtering steps. The genotypes on chrY cannot be directly compared with the Platinum genotypes because chrY calls were not provided in the VCF file of the Platinum individuals. To assess the performance of chrY genotyping, we compared our results to the genotypes of 11 men from the HGDP panel^25^ (http://cdna.eva.mpg.de/denisova/). The overall concordance of the haploid genotype calls of FastGT and the genotype calls in these VCF files was 99.97%. The fraction of non-canonical genotypes (no-calls) in the FastGT output was 1.27% (Table S3).

We also tested the concordance of chrY genotypes in seven father-son pairs in CEPH pedigree 1463 (http://www.ebi.ac.uk/ena/data/view/ERP001960). We assume that changes in chrY genotypes should not occur within one generation. Only one marker (rs199503278) showed conflicting genotypes in any of these father-son pairs. A visual inspection revealed problems with the reference genome assembly in this region, which resulted in conflicting *k*-mer counts and conflicting genotypes from different *k*-mer pairs of the same SNV. This marker was removed from the dataset because it had a high likelihood of causing similar problems in other individuals.

### Effect of genome coverage on FastGT performance

We also studied how the genome sequencing depth affects the performance of FastGT. The Platinum genomes have a coverage depth of approximately 50x, but in most study scenarios, sequencing to a lower coverage is preferred because it optimizes costs. For this analysis, we compiled different-sized subsets of FASTQ sequences from the Platinum individual NA12878 and measured the concordance between called genotypes and genotypes from the Platinum dataset. We observed that the concordance rate of non-reference genotypes (AB and BB) declines significantly as the coverage drops below 20x (Figure 4).

**Figure 4.**
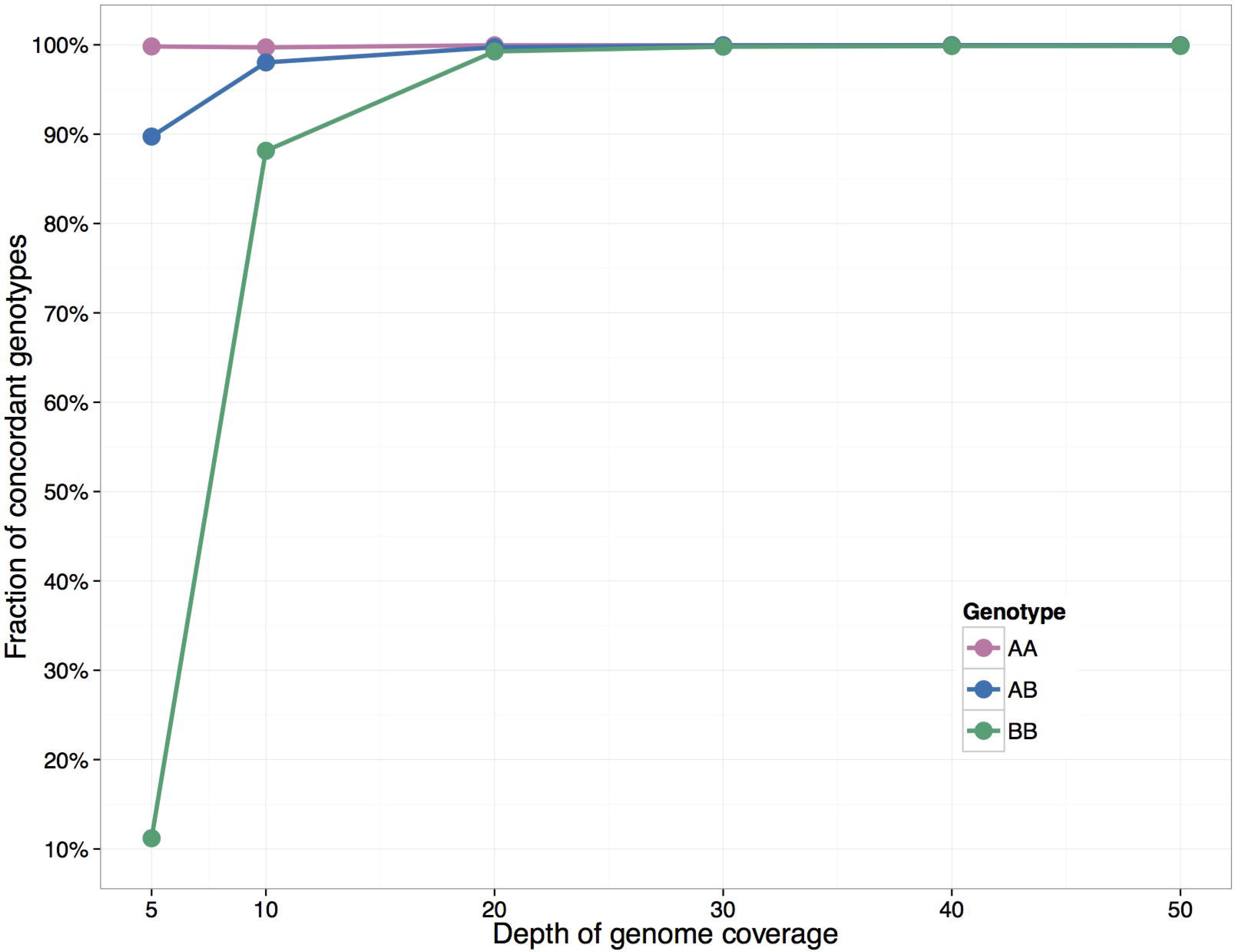
Effect of genome coverage on the concordance of genotypes. The accuracy of calling non-reference variants starts to decline as the genome coverage drops below 20x. Only the accuracy of the non-reference allele (genotypes AB and BB) calls declines significantly as the coverage drops because the higher prior probability of the reference allele has a stronger influence on the final decision of the Bayesian classifier in situations where the coverage is low (which increases the bias toward the more common allele).

### Relationship between *k*-mer length and number of usable variants

An obvious question is how the *k*-mer length affects the performance of FastGT. We used 25 nucleotides long *k*-mers throughout this article, but FastGT is able to use other *k*-mer lengths between 16 and 32 as well. We tested how many markers from dbSNP would remain usable for FastGT at different values of *k*. From 47 millions validated markers 7-17% markers are removed in filtering step 1 due to closely located SNVs (Figure 5). In filtering step 2 markers are removed if they have no unique *k*-mer pairs in the expanded reference genome. As expected, a rather large number of markers are eliminated from the dataset if *k*-mers shorter than 20 nucleotides are used. However, the number of usable markers does not increase significantly for *k* larger than 24. Thus, *k* values between 24 and 32 should be equally suitable for analyzing the human genome with FastGT. We have not compared the accuracy of genotype calls of different *k*-mer lengths. However, we expect it to be relatively independent of *k*-mer length. Two main factors that might influence the accuracy (concordance) of genotypes are non-specific counts from shorter *k*-mers and drop of effective coverage of *k*-mers. The effective coverage is negatively correlated with *k*-mer length. This negative correlation is caused by the higher chance of accumulating sequencing errors within longer *k*-mers and by the end effects of the reads (lower number of long *k*-mers per sequencing read). On the other hand, shorter *k*-mers are more likely to pick up non-specific sequences due to sequencing errors and unknown variations in human genomes. Overall, these effects influence the effective coverage of *k*-mers and are only critical if genome coverage is low or if *k*-mer is shorter than 20 nucleotides. At high coverage (>20x) conditions the *k*-mer length should not have significant influence to genotype accuracy.

**Figure 5.**
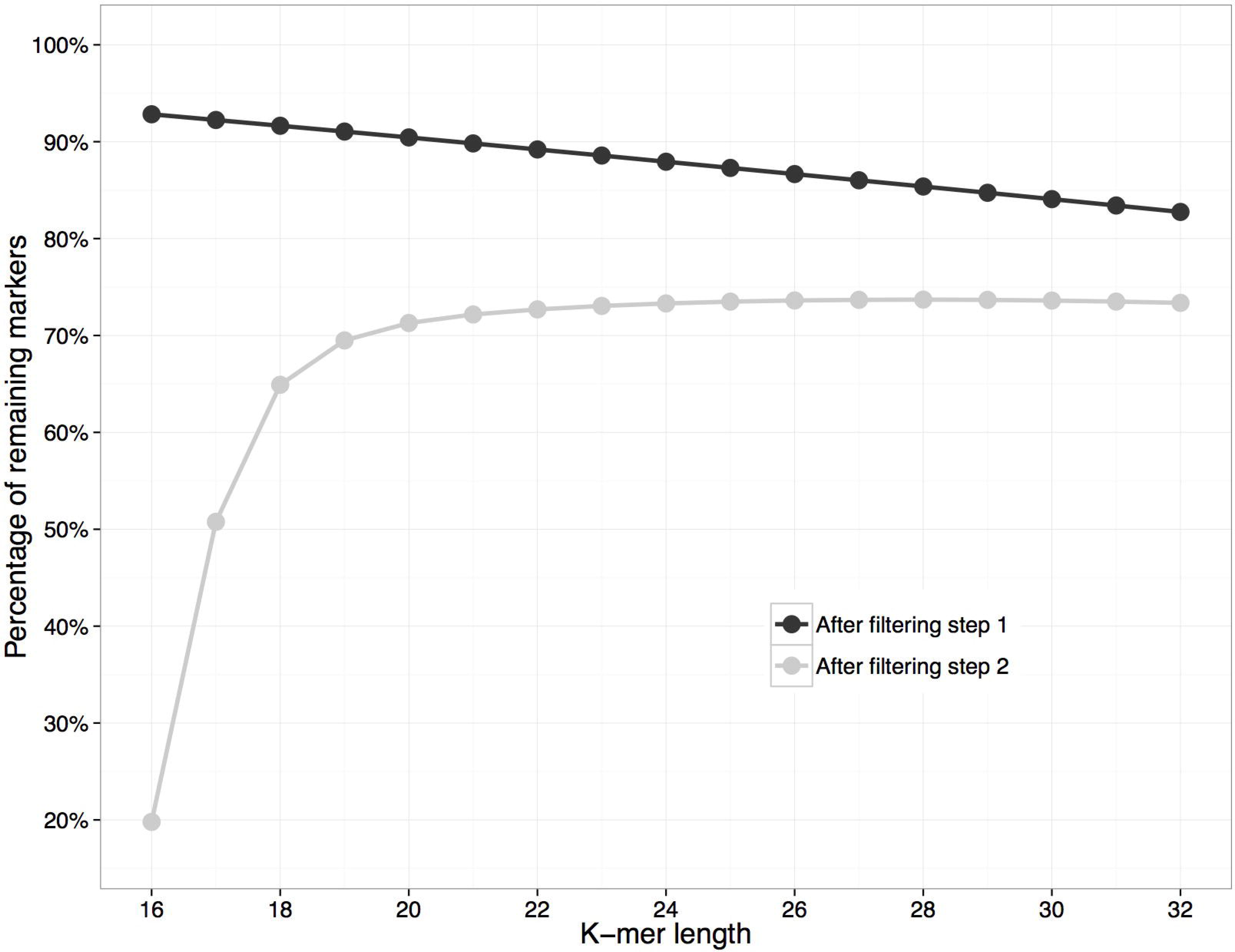
Effect of *k*-mer length on filtering SNV markers. Lines show the fraction of remaining markers after filtering steps 1 and 2. The step 1 removes SNVs that have other marker within *k* nucleotides on both sides. The step 2 removes SNVs that have no unique *k*-mers in the expanded reference genome. Y-axis indicates the fraction of remaining markers after each filtering step. 100% in this figure corresponds to 46,954,719 SNVs from the dbSNP that were fed to filtering pipeline.

### Time and memory usage

The entire process of detecting 30 million SNV genotypes from the sequencing data of a single individual (30x coverage, 2 FASTQ files, 115GB each) takes approximately 40 minutes on a server with 32 CPU cores. Most of this time is allocated to counting *k*-mer frequencies by 

~~~
gmer_counter
~~~

. The running time of 

~~~
gmer_counter
~~~

 is proportional to the size of the FASTQ files because the speed-limiting step of 

~~~
gmer_counter
~~~

 is reading the sequence data from a FASTQ file. However, the running time is also dependent on the number of FASTQ files (Figure 6) because simultaneously reading from multiple files is faster than processing a single file. Genotype calling with 

~~~
gmer_caller
~~~

 takes approximately 2-3minutes with 16 CPU cores.

**Figure 6.**
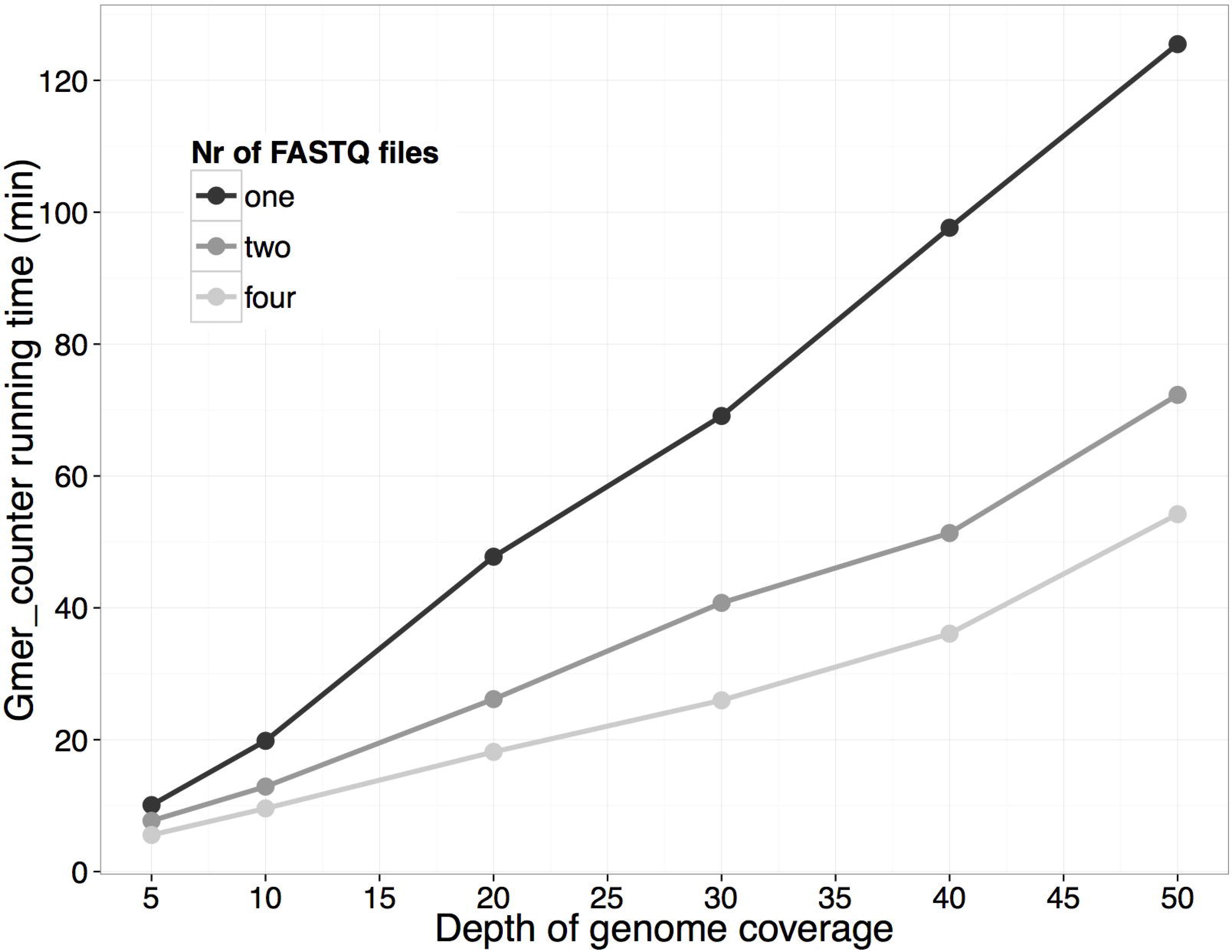
The time spent counting *k*-mer frequencies is proportional to the genome coverage (because of the larger FASTQ files). 

~~~
Gmer_counter
~~~

 is able to read data from multiple files simultaneously; thus, it runs faster if the sequence data are distributed between different files (e.g., files with paired reads).

The minimum amount of required RAM is determined by the size of the data structure stored in memory by 

~~~
gmer_counter
~~~

. We have tested 

~~~
gmer_counter
~~~

 on Linux computer with 8 GB of RAM. However, server-grade hardware (multiple CPU cores and multiple fast hard drives in RAID) is required to achieve the full speed of 

~~~
gmer_counter
~~~

 and 

~~~
gmer_caller
~~~

.

## METHODS

### Compilation of database of unique *k*-mers

A *k*-mer length of 25 was used throughout this study, and the *k*-mers for genotyping were selected by the following filtering process (see also Figure S4). First, the validated single nucleotide variants (SNVs), as well as the validated and common indels, were extracted from the dbSNP database build 146^26,27^. Indels were used for testing the uniqueness of *k*-mers only; they are not included in the database of variants. For every bi-allelic SNV from this set, two sequences surrounding this SNV location were created: the sequence of the human reference genome (GRCh37) and the sequence variant corresponding to the alternative allele. The sequences were shortened to eliminate any possible overlap with neighboring SNVs or common indels. Essentially, this filtering step removed all of the SNVs that were located between two other SNVs (or indels) with less than 25bp between them. This step was chosen to avoid the additional complexity of counting and calling algorithms because of the multiple combinations of neighboring SNV alleles. For all these SNVs that had variant-free sequences of at least 25bp, the sequences were divided into 25-mer pairs.

In the second filtering step, we tested the uniqueness of the 25-mers compiled in the previous step. The uniqueness parameter was tested against the “expanded reference genome,” which is a set of 25-mers from the reference genome plus all possible alternative 25-mers containing the non-reference alleles of the SNVs and indels. A *k*-mer pair is considered unique if both *k*-mers occur no more than once in the “expanded reference genome”. All non-unique *k*-mer pairs were removed from the list. The 

~~~
Glistcompare
~~~

 tool^28^, which performs set operations with sorted *k*-mer lists, was used in this step. The *k*-mer pairs demonstrating uniqueness even with one mismatch were preferred. This constraint was added to reduce the risk of forming an identical *k*-mer by a rare point mutation or a sequencing error.

In the third step, the *k*-mers were further refined using the *k*-mer frequencies and genotypes in a set of sequenced individual genomes. For this purpose, the *k*-mer counts and genotypes were calculated for all SNVs of 50 random individuals whose DNA was collected and sequenced during the Center of Translational Genomics project at the University of Tartu. Twenty-five men and 25 women were used for filtering the autosomal SNVs; for chrX and chrY, 50 men were used. The sequencing depth in these individuals varied between 21 and 45. Three different criteria were used for removing k-mer pairs and SNVs in this step. First, we excluded all chrY markers that had *k*-mer frequency higher than 3 in more than one woman. Second, autosomal *k*-mers showing abnormally high frequencies (greater than 3 times the median count) in more than one individual were removed. Third criterion was based on unexpected genotypes: the SNVs that produced a non-canonical allele count in more than one individual out of 50 were removed from the dataset. The non-canonical allele count is any value other than two alleles in autosomes or a single allele in male chrX and chrY. The criteria used in filtering step 3 should remove SNVs located in the regions that are unique in the reference genome, but frequently duplicated or deleted in real individuals.

The final set contained 30,238,283 SNVs usable by FastGT, with 6.8% (2,063,839) located in protein-coding regions. A detailed description of the filtering steps used in this article is shown in in Figure S5. The number of markers removed in each step is shown in Table S1.

### Statistical framework

The statistical framework for Empirical Bayes Classifier implemented in 

~~~
gmer_caller
~~~

 is described in Supplementary File.

### Generating and analyzing simulated data

FastGT was tested on simulated reads. Simulated sequencing reads were generated using WgSim (version 0.3.1-r13) software from samtools package^3^. The following parameters were used: base_error_rate=0.005, outer_distance_between_the_two_ends=500, standard_deviation=50, length_of_the_first_read=100, length_of_the_second_read=100. We used the base error rate 0.005 (0.5%) because this is similar to error rate typically observed in Illumina HiSeq sequencing data. We estimated average error rate in the raw reads of high-coverage genomes from Estonian Genome Center by counting the fraction of erroneous *k*-mers. The error rates in 100 individuals varied between 0.0030 and 0.0082, with average 0.0048 (CI95%=0.0002). Previous studies have reported similar overall error rate in raw reads generated by Illumina HiSeq, varying between 0.002 and 0.004,^29,30^. The sequencing reads were simulated with different coverages: about 5, 10, 20, 30 and 40. The number of read pairs generated were 80 million, 160 million, 320 million, 480 million and 640 million respectively. Reads were generated from standard reference genome, version GRCh37. For Figure 3 the reads were also simulated using real SNV information for 5 individuals from 5 different populations (CEU, CHS, YRI, PUR and EST). The following individuals were used in simulations: HG00512 (CHS), NA19238 (YRI), HG00731 (PUR), NA12877 (CEU) and V00055 (EST). Their sequencing data was retrieved from 1000G project repository at ftp://ftp.1000genomes.ebi.ac.uk/vol1/ftp/data_collections/hgsv_sv_discovery/data/ (CHS, YRI and PUR), from the ftp://ftp.sra.ebi.ac.uk/vol1/ERA172/ERA172924/bam/ (CEU) or from the Estonian Genome Center. For each of these individuals, the standard reference genome was used as base and the corresponding SNV genotypes from their VCF files were added to generate the reads with realistic variants. The SNV genotypes were calculated from BAM or CRAM files using Genome Analysis Toolkit version 3.6^4^.

The sensitivity and specificity were calculated for each individual and for each coverage. True positive was defined as AB or BB genotype that was correctly called by FastGT in simulated data. True negative values were defined as correctly called AA genotypes. The genotypes called from sex chromosomes were not used for sensitivity and specificity calculations.

### Testing genotype concordance

Version 20160503 of the FastGT package was used throughout this study. For the concordance analysis with the Platinum genotypes, 

~~~
gmer_counter
~~~

 and 

~~~
gmer_caller
~~~

 were run with the default options. The performance was tested on a Linux server with 32 CPU cores, 512GB RAM, and IBM 6Gbps and SAS 7200rpm disk drives in a RAID10 configuration.

High-quality genotypes were retrieved from the Illumina Platinum Genomes FTP site at ftp://ussd-ftp.illumina.com/hg38/2.0.1/.

BAM-format files of NA12877 and 12878 were downloaded from ftp://ftp.sra.ebi.ac.uk/vol1/ERA172/ERA172924/bam/NA12877_S1.bam and ftp://ftp.sra.ebi.ac.uk/vol1/ERA172/ERA172924/bam/NA12878_S1.bam.

FASTQ files were downloaded from the European Nucleotide Archive at http://www.ebi.ac.uk/ena/data/view/ERP001960. FASTQ files for the chrY genotype comparison were created from the corresponding BAM files using SAMtools bam2fq version 0.1.18. The read length of the Platinum genomes was 101 nucleotides.

Illumina HumanOmniExpress microarray genotypes and Illumina NGS data (read length 151 nt) for individuals V00278, V00328, V00352, V00369, V00402, V08949, V09325, V09348, V09365, and V09381 were obtained from the Estonian Genome Center. For the concordance analysis with the microarray genotypes, 

~~~
gmer_caller
~~~

 was run with the microarray markers (504,173) only.

The 5x, 10x 20x, 30x, and 40x data points for Figure 4 were created using random subsets of reads from raw FASTQ files of 50x coverage from the Platinum individual NA12878.

### Code availability

The binaries of FastGT package and *k*-mer databases described in the current paper are available on our website, http://bioinfo.ut.ee/FastGT/. The source code is available at GitHub (https://github.com/bioinfo-ut/GenomeTester4/). 

~~~
Gmer_counter
~~~

 and 

~~~
gmer_caller
~~~

 are distributed under the terms of GNU GPL v3, and the *k*-mer databases are distributed under the Creative Commons CC BY-NC-SA license.

## DISCUSSION

FastGT is a flexible software package that performs rapid genotyping of a subset of previously known variants without a loss of accuracy. Another similar approach of genotype calling has been published before^24^. Both methods need to pre-process the reference genome and personal short-read data. Our method pre-processes the genome by selecting the SNVs and compiling the database of *k*-mers that can be used for calling these SNVs. The short-read data is pre-processed by counting and storing the *k*-mer frequencies using 

~~~
gmer_counter
~~~

. The method by Kimura and Koike uses dictionary-based approach for storing both reference sequence and short reads. The dictionary is implemented by means of the Burrows-Wheeler transform (BWT). The main advantage of BWT is the ability of storing and comparing long strings efficiently. Therefore, this method can be used to call all SNVs, including those that are in repeated genomic regions. FastGT uses fixed length *k*-mer with maximum length of 32. This limits the number of variants that can be called from the human genome. On the other hand, using fixed length k-mers allows faster processing of data due to 64-bit architecture of computer hardware. Thus, FastGT essentially sacrifices calling some SNVs (up to 36%) from difficult genomic regions to minimize data processing time. Another difference between FastGT and the method used by Kimura and Koike is handling of the *de novo* mutations. Kimura and Koike implemented two methods (drop-scan and step-scan) to detect de novo variants based on k-mer coverage and/or by local alignment of surrounding region. FastGT has currently no ability to call *de novo* variants and is limited to calling sub-sets of pre-defined variants. Thus, FastGT functions in principle as a large digital microarray with millions of probes.

Numerous software packages can organize the raw sequencing data of each individual into comprehensive *k*-mer lists^28,31–34^, which can be later used for fast retrieval of *k*-mer counts. However, the compilation of full-genome lists is somewhat inefficient if the lists are only used once and then immediately deleted. FastGT uses adaptive radix tree, which allows us to store frequencies for only the *k*-mers of interest, instead of for all *k*-mers from the genome. This approach is particularly useful for genotyping only a small number of variants from each individual. Storing only the frequencies of relevant k-mers avoids the so-called “curse of deep sequencing,” in which a higher coverage genome can overwhelm the memory or disk requirements of the software^35^. The disk and memory requirements of FastGT are not directly affected by the coverage of sequencing data.

Our analysis focuses on genotyping SNVs. However, FastGT is not limited to identifying SNVs. Any known variant that can be associated with a unique and variant-specific *k*-mer can be detected with FastGT. For example, short indels could be easily detected by using pairs of indel-specific *k*-mers. In principle, large indels, pseudogene insertions, polymorphic Alu-elements, and other structural variants could also be detected by *k*-mer pairs designed over the breakpoints. However, the detection of structural variants relies on the assumption that these variants are stable in the genome and have the same breakpoint sequences in all individuals, which is not always true for large structural variants. The applicability of FastGT for detecting structural variants requires further investigation and testing.

This software has only been used with Illumina sequencing data, which raises the question of whether our direct genotyping algorithm is usable with other sequencing technologies. In principle, *k*-mer counting should work with most sequencing platforms that produce contiguous sequences of at least *k* nucleotides. The uniformity of coverage and the fraction of sequencing errors in raw data are the main factors that influence *k*-mer counting because a higher error rate reduces the number of usable *k*-mers and introduces unwanted noise. The type of error is less relevant because both indel-type and substitution-type errors are equally deleterious for *k*-mer counting.

NGS data are usually stored in BAM format, and the original FASTQ files are not retained. In this case, the FASTQ file can be created from available BAM files. This can be performed by a number of software packages (Picard, bam2fq from SAMtools package^1^, bam2fastx from TopHat package^36^). We have tested FastGT software with raw FASTQ files and FASTQ files generated from the BAM-formatted files and did not observe significant differences in the *k*-mer counts or genotype calls. In principle, care should be taken to avoid multiple occurrences of the same reads in the resulting FASTQ file. Regardless of the method of genome analysis, contamination-free starting material, diligent sample preparation, and sufficient genome coverage are the ultimate pre-requisites for reliable results. The “garbage in, garbage out” principle applies similarly to mapping-based genome analyses and *k*-mer based genome analyses.

## ACKNOWLEDGEMENTS

This work was funded by institutional grant IUT34-11 from the Estonian Ministry of Education and Research, grant SP1GVARENG from the University of Tartu, and the EU ERDF grant No. 2014-2020.4.01.15-0012 (Estonian Center of Excellence in Genomics and Translational Medicine). The cost of the NGS sequencing of the individuals from the Estonian Genome Center was partly covered by the Broad Institute (MA, USA) and the PerMed I project from the TERVE program. The genome data was collected and used with ethical approval Nr. 206T4 (obtained for the project SP1GVARENG). The computational costs were partly covered by the High Performance Computing Centre at the University of Tartu. The authors thank Märt Roosaare, Ulvi Talas, and Priit Palta for performing a critical reading of the manuscript.

## AUTHOR CONTRIBUTIONS

FDP compiled the databases of unique *k*-mers and conducted the genotype concordance analyses. LK invented the data structures and algorithms for 

~~~
gmer_counter
~~~

 and 

~~~
gmer_caller
~~~

 and implemented their code in C. MM wrote the initial code of the Bayesian classifier for genotype calling and supervised the development of a statistical framework. TP validated the genotyping results by performing a manual analysis of BAM files and providing expertise for NGS data management. ML performed an initial survey of the optimal number of *k*-mer pairs per variant. MR supervised the work and wrote the final version of the manuscript.

## REFERENCES

1. Eberle, M. A. et al. A reference dataset of 5.4 million phased human variants validated by genetic inheritance from sequencing a three-generation 17-member pedigree. bioRxiv (2016).

2. Li, H. & Durbin, R. Fast and accurate long-read alignment with Burrows-Wheeler transform. Bioinformatics 26, 589–95 (2010).

3. Li, H. et al. The Sequence Alignment/Map format and SAMtools. Bioinformatics 25, 2078– 2079 (2009).

4. McKenna, A. et al. The Genome Analysis Toolkit: a MapReduce framework for analyzing next-generation DNA sequencing data. Genome Res. 20, 1297–303 (2010).

5. Langmead, B. & Salzberg, S. L. Fast gapped-read alignment with Bowtie 2. Nat. Methods 9, 357–9 (2012).

6. Highnam, G. et al. An analytical framework for optimizing variant discovery from personal genomes. Nat. Commun. 6, 6275 (2015).

7. Zook, J. M. et al. Integrating human sequence data sets provides a resource of benchmark SNP and indel genotype calls. Nat. Biotechnol. 32, 246–251 (2014).

8. O’Rawe, J. et al. Low concordance of multiple variant-calling pipelines: practical implications for exome and genome sequencing. Genome Med. 5, 28 (2013).

9. Pirooznia, M. et al. Validation and assessment of variant calling pipelines for next-generation sequencing. Hum. Genomics 8, 14 (2014).

10. Li, H. Toward better understanding of artifacts in variant calling from high-coverage samples. Bioinformatics 30, 2843–51 (2014).

11. Derrien, T. et al. Fast computation and applications of genome mappability. PLoS One 7, (2012).

12. Lee, H. & Schatz, M. C. Genomic dark matter: The reliability of short read mapping illustrated by the genome mappability score. Bioinformatics 28, 2097–2105 (2012).

13. Weisenfeld, N. I. et al. Comprehensive variation discovery in single human genomes. Nat. Genet. 46, 1350–5 (2014).

14. Wen, J., Chan, R. H. F., Yau, S.-C., He, R. L. & Yau, S. S. T. K-mer natural vector and its application to the phylogenetic analysis of genetic sequences. Gene 546, 25–34 (2014).

15. Ondov, B. D. et al. Mash: fast genome and metagenome distance estimation using MinHash. (2015). doi:10.1101/029827

16. Haubold, B., Klötzl, F. & Pfaffelhuber, P. andi: fast and accurate estimation of evolutionary distances between closely related genomes. Bioinformatics 31, 1169–75 (2015).

17. Hasman, H. et al. Rapid whole-genome sequencing for detection and characterization of microorganisms directly from clinical samples. J. Clin. Microbiol. 52, 139–46 (2014).

18. Wood, D. E. & Salzberg, S. L. Kraken: ultrafast metagenomic sequence classification using exact alignments. Genome Biol. 15, R46 (2014).

19. Roosaare, M. et al. StrainSeeker: fast identification of bacterial strains from unassembled sequencing reads using user-provided guide trees. (2016). doi:10.1101/040261

20. Song, L., Florea, L. & Langmead, B. Lighter: fast and memory-efficient sequencing error correction without counting. Genome Biol. 15, 509 (2014).

21. Marçais, G., Yorke, J. A. & Zimin, A. QuorUM: An Error Corrector for Illumina Reads. PLoS One 10, e0130821 (2015).

22. Lim, E.-C. et al. Trowel: a fast and accurate error correction module for Illumina sequencing reads. Bioinformatics 30, 3264–5 (2014).

23. Zhao, X. et al. EDAR: an efficient error detection and removal algorithm for next generation sequencing data. J. Comput. Biol. 17, 1549–60 (2010).

24. Kimura, K. & Koike, A. Ultrafast SNP analysis using the Burrows-Wheeler transform of short-read data. Bioinformatics 31, 1577–83 (2015).

25. Meyer, M. et al. A high-coverage genome sequence from an archaic Denisovan individual. Science 338, 222–6 (2012).

26. NCBI Resource Coordinators. Database resources of the National Center for Biotechnology Information. Nucleic Acids Res. 44, D7–19 (2016).

27. Sherry, S. T., Ward, M. & Sirotkin, K. dbSNP-database for single nucleotide polymorphisms and other classes of minor genetic variation. Genome Res. 9, 677–9 (1999).

28. Kaplinski, L., Lepamets, M. & Remm, M. GenomeTester4: a toolkit for performing basic set operations - union, intersection and complement on k-mer lists. Gigascience 4, 58 (2015).

29. Ross, M. G. et al. Characterizing and measuring bias in sequence data. Genome Biol. 14, R51 (2013).

30. Schirmer, M., D’Amore, R., Ijaz, U. Z., Hall, N. & Quince, C. Illumina error profiles: resolving fine-scale variation in metagenomic sequencing data. BMC Bioinformatics 17, 125 (2016).

31. Marçais, G. & Kingsford, C. A fast, lock-free approach for efficient parallel counting of occurrences of k-mers. Bioinformatics 27, 764–770 (2011).

32. Deorowicz, S., Kokot, M., Grabowski, S. & Debudaj-Grabysz, A. KMC 2: Fast and resource-frugal k-mer counting. Bioinformatics 31, 1569–1576 (2014).

33. Rizk, G., Lavenier, D. & Chikhi, R. DSK: K-mer counting with very low memory usage. Bioinformatics 29, 652–653 (2013).

34. Roy, R. S., Bhattacharya, D. & Schliep, A. Turtle: Identifying frequent k-mers with cache-efficient algorithms. Bioinformatics 30, 1950–1957 (2014).

35. Roberts, A. & Pachter, L. RNA-Seq and find: entering the RNA deep field. Genome Med. 3, 74 (2011).

36. Trapnell, C., Pachter, L. & Salzberg, S. L. TopHat: Discovering splice junctions with RNA-Seq. Bioinformatics 25, 1105–1111 (2009).

